# Quinpirole ameliorates the dysfunction of microglia in human LRRK2-R1441G transgenic mice

**DOI:** 10.1101/2025.02.23.639776

**Authors:** Yuanxin Chen, Shiquan Cui, Huaixin Wang, Chenbo Zeng, Hui Zhang

## Abstract

Microglia-mediated neuroinflammation is a key contributor to Parkinson’s disease (PD) pathogenesis. Leucine-rich repeat kinase 2 (LRRK2), the leading genetic contributor to both familial and sporadic PD, has been implicated in driving this connection. However, its precise role remains incompletely understood due to technical challenges. To address this, we utilized a bacterial artificial chromosome (BAC) transgenic mouse model overexpressing human LRRK2-R1441G, which replicates key features of PD. These mice were crossed with Cx3cr1–EGFP mice to enable assessment of microglial dynamics and function using two-photon imaging in awake mice *in vivo* and acute brain slices *ex vivo*. Furthermore, spatial transcriptomic analysis was performed using GeoMx Digital Spatial Profiler technology to compare transgenic mice with their wild-type counterparts. The R1441G mutation upregulated antigen processing and presentation pathways, increased activated microglia, and enhanced microglial polarization in the dorsal striatum. Mutant microglia exhibited reduced motility and slower responses to focal injury, with processes retracting faster and extending more slowly. Quinpirole, a dopamine D2 receptor (D2R) agonist, successfully reversed microglial deficits. This study provides the first evidence that pathogenic LRRK2 mutations alter microglial motility and responsiveness *in vivo*, highlighting D2R activation as a promising therapeutic strategy to mitigate neuroinflammation and neurodegeneration in PD.

## Introduction

Parkinson’s disease (PD), the second most prevalent neurodegenerative disorder, is characterized by the progressive degeneration of dopaminergic neurons in the substantia nigra pars compacta (SNc) and a subsequent reduction in dopamine release within the dorsal striatum (dSTR). Emerging evidence highlights the critical role of inflammatory processes in PD pathogenesis^1^, with microglial activation linked to immune dysregulation and neuroinflammation, factors closely associated with aging and neurodegenerative diseases.^2^

To date, approximately 20 genes have been implicated in the pathogenesis of PD, with mutations in LRRK2 being the most identified. Li *et al*.^3^ reported motor deficits and reduced dopamine release in the LRRK2 hR1441G animal model. Some studies have suggested that LRRK2 plays a key role in linking immune function with PD susceptibility, given its elevated expression in immune cells, including dendritic cells and macrophages.^4^ Microglia, the resident brain macrophages, exhibit rapid response to brain injury and protect the brain from further injury^5^. Additionally, research indicates that microglia lacking LRRK2 exhibit increased motility compared to control cells, suggesting that LRRK2 acts as a negative regulator of microglial motility.^6^

Dopamine, a critical neurotransmitter for motor, emotional, and cognitive functions in the central nervous system (CNS), exerts immunomodulatory effects on immune cells.^7^ The dopamine receptor D2 receptor (D2R) is expressed in activated resident microglia, and D2R agonists enhances the responsiveness of cultured microglia to pro-inflammatory stimuli.^8^ Notably, aging in the human brain is associated with a decline in D2R density in both the striatum and extracerebral regions, correlating with progressive cognitive and motor function impairments in the elderly.^9^ Shao *et al* and Zhang *et al*^10,11^ reported that D2R activation in glial cells, including astrocytes and microglia, through treatment with the agonist quinpirole, typically suppresses neuroinflammation in the CNS via mechanisms involving αB-crystallin.

Microglia are highly dynamic cells, using their branched processes to continuously monitor the brain for abnormalities and injuries.^12,13^ Understanding how LRRK2 mutations influence microglial activity and neuroinflammation requires studying these cells within their native environment over time. However, due to technical challenges, most research to date has relied on cultured or isolated microglia, which alters their natural properties and limits insights into their role within intact brain microenvironments. Therefore, the impact of pathogenic LRRK2 mutations on microglial dynamics and the potential of D2R agonists to mitigate LRRK2-related neuroinflammation remain poorly understood. In the present study, using time-lapse two-photon imaging in awake mice and acute brain slices, we aimed to address two key questions: (1) to determine whether the LRRK2 hR1441G mutation drives neuroinflammation through microglial polarization and disrupts microglial motility, and (2) to evaluate whether D2 receptor agonists can reverse the functional impairments in microglia caused by the pathogenic LRRK2 mutation.

## Materials and methods

### Animals and brain slice preparation

Animal procedures complied with NIH guidelines and received IACUC approval from Thomas Jefferson University and The University of Georgia. BAC LRRK2 (hR1441G) transgenic mice were obtained from Weill Medical College of Cornell University (FVB/N background, Jackson Laboratory). CX3CR1-GFP mice (Jackson Laboratory) were backcrossed onto FVB/N for eight generations and crossed with hR1441G mice to produce GFP-expressing hR1441G and non-transgenic (nTg) lines. All mice were on FVB/N mouse background (Taconic).

Anesthetized mice were perfused with ice-cold, oxygenated cutting solution, and 300 µm coronal slices were obtained and incubated in oxygenated aCSF at 34°C for 30-45 minutes, before being transferred to room temperature.

### Tissue preparation and immunofluorescence staining

Animals were anesthetized via intraperitoneal injection of a ketamine/xylazine mixture (100 mg/10 mg/kg) and subsequently underwent transcardial perfusion with 4% paraformaldehyde (PFA) fixative. Fixed brains were coronally sectioned at a thickness of 30 μm using a cryostat microtome. Immunostaining was performed with primary antibodies (anti-TH, anti-IBA1, anti-CD68) and corresponding fluorescently conjugated secondary antibodies. Imaging was conducted on a Zeiss LSM 900 confocal microscope.

### Cranial window surgery: polished and reinforced thinned skull (PoRTS) procedure

Three to four male pairs of hR1441G-Tg/CX3CR1-GFP^+^ and hR1441G-WT (nTg)/ CX3CR1-GFP^+^ mice (6–7 months old) underwent PoRTS surgery, following the protocol described by

Drew *et al*.^14^ Experiments were performed only if physiological parameters were maintained within normal ranges.

### Two-photon laser ablation

Laser-induced injury was generated by directing a two-photon laser beam to a confined region at a depth of 70–150 µm through a thinned, intact skull for *in vivo* imaging, or at a depth of 30–50 µm below the surface of brain slices in the dorsal striatum for *ex vivo* imaging.^13^

### 2PLSM: imaging of microglia in acute brain slices and *in vivo*

The upright laser scanning microscope (BX61WI, Olympus), coupled with a Ti: sapphire pulsed laser system (80 MHz repetition rate, <100 fs pulse width, Coherent Inc.), and controlled using Prairie View 5.3 software (Bruker), was employed for two-photon fluorescence imaging (Supplementary Fig. 1) for *in vivo* and *ex vivo* imaging of microglia.

### Quinpirole treatment

Brain slices were treated with 3 µM quinpirole, and in *vivo* imaging was conducted following a single intraperitoneal injection of quinpirole (0.5 mg/kg).^15^

### Imaging data analysis

Images were processed using Fiji (NIH) and Matlab (Version 8.5.0 R2015a, MathWorks). To analyze the dynamics of microglial protrusions and retractions, the length of microglial processes and the velocity of length changes were assessed from maximum intensity projection (MIP) as described by Nimmerjahn *et al*^16^ (Supplementary Fig. 2). Additionally, microglial response to laser ablation were quantified following the methodology described by Davalos.^13^ Further detailed methods are provided in the supplementary materials.

### Digital spatial profiling

Formalin-fixed paraffin-embedded (FFPE) sections were processed and subjected to GeoMx RNA profiling (Nanostring Technologies, Seattle, WA, USA). ROIs/AOIs were defined using TH and IBA1 markers, and gene set enrichment analysis (GSEA) was performed to identify enriched pathways in LRRK2 mutant microglia. Further detailed methods are provided in the supplementary materials.

### Statistical analysis

Data are presented as mean ± standard deviation (SD) where applicable. Statistical analysis was performed with GraphPad Prism 8 (GraphPad software Inc.). Unpaired Mann-Whitney test, one-way ANOV, and two-way ANOVA, were applied, with a P-value of < 0.05 considered statistically significant.

## Results

### LRRK2 hR1441G mutation caused neuroinflammation and microglia polarization

To examine whether microglia with LRRK2 hR1441G mutation is activated, GeoMx digital spatial profiler (DSP) analysis was performed on microglia in the SNc, comparing gene expression profiles between hR144G TG and WT mice. The analysis revealed 18 significantly upregulated and 27 downregulated genes in the mutant microglia (Fig. 1A-B and Supplementary Fig. 3A-B). The top 20 significantly enrich terms among the upregulated genes were identified, with most of them associated with neuroinflammation, such as “Antigen processing and presentation of exogenous peptide antigen via MHC class”, “mast cell activation” and “steroid catabolic process” (Fig. 1C-E). Additionally, significant neuroinflammation-related enrichment terms among the downregulated genes, such as “tryptophan transport” and “cell communication by electrical coupling,” were found to be downregulated. These findings suggest that hR1441G mutation triggers microglia activation and induces neuroinflammation.

**Figure 1.**
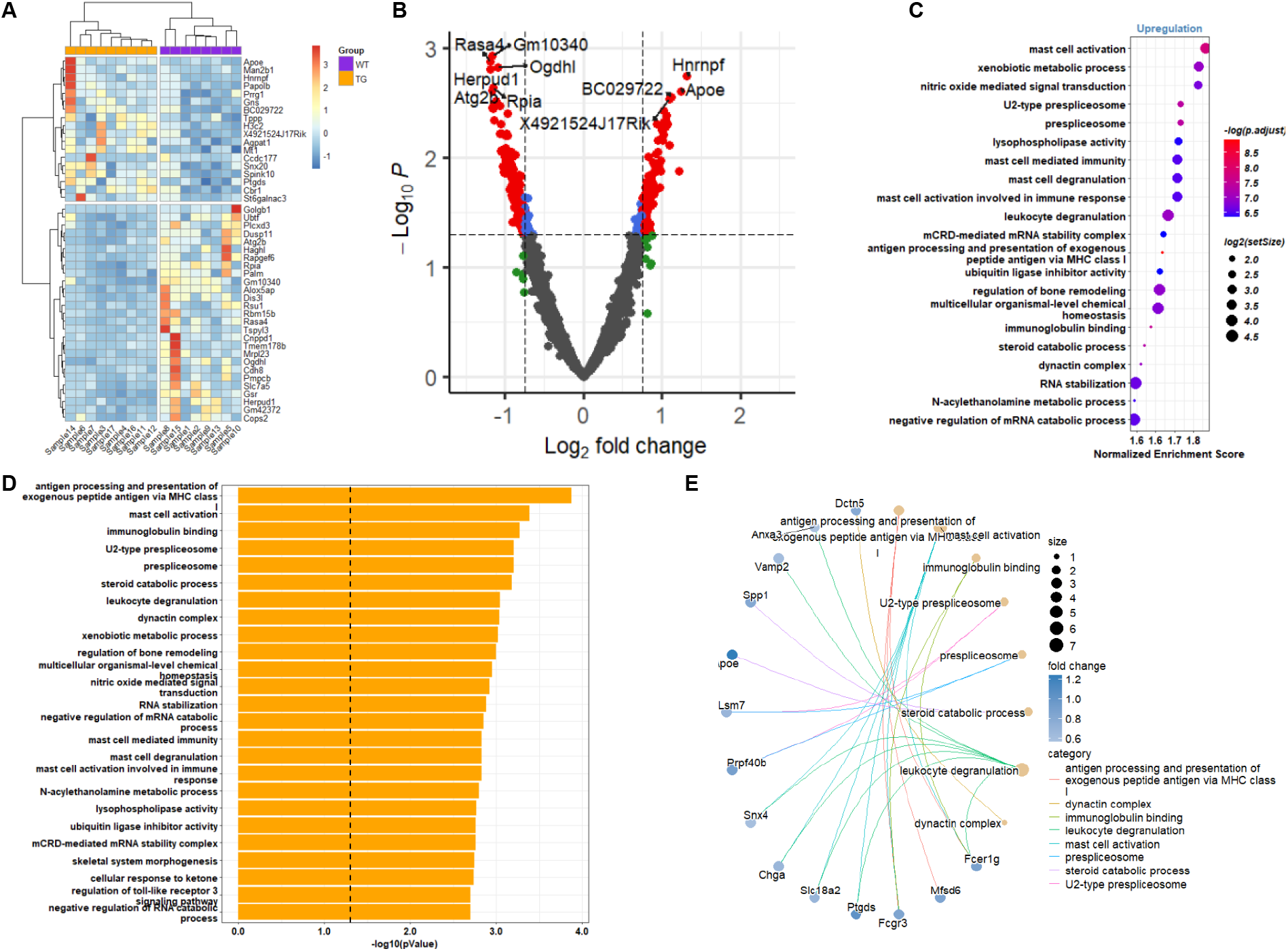
GeoMx digital spatial profiling (DSP) of microglia indicates microglial neuroinflammation. **(A)** Heatmap displaying significantly differentially expressed genes. **(B)** Volcano plot illustrating the distribution of differentially expressed genes. **(C)** Gene ontology enrichment analysis of upregulated genes, with terms colored by p-values and sized according to gene set size. **(D)** Bar plot showing the top 20 enriched terms, ordered by p-value. **(E)** Gene network visualization of enriched terms.

To determine whether the hR1441G mutation induces microglial polarization in the dSTR, IBA1 and TH staining were employed to assess the distribution of microglia in hR1441G transgenic (TG) mice.^3^ As shown in Fig. 2A1-A2, B1-B2, C1-C2 and E, no significant difference in microglia coverage was observed between WT and TG mice at 1-month-old age (WT, N = 3; TG, N = 4; P = 0.4893). However, at 5 months of age, TG mice exhibited a significantly higher microglia coverage area compared to WT littermates (4 pairs; P = 0.0149), although both groups exhibited a decrease compared to 1-month-old mice. At 22 months of age, TG mice displayed markedly higher microglia coverage than WT mice (3 pairs, P < 0.0001), with both groups exhibiting increased microglial coverage compared to the 5-month-old cohort. These findings are consistent with the lifespan dynamics of microglia reported by Nikodemova *et al*.^17^ where the numbers declined from 1 month old to young adulthood. The rate of decline was slower in TG mice (Fig. 2E), suggesting that neuroinflammation may drive microglial proliferation and monocytes differentiation in the brain.^18^ To further explore microglial polarization, CD68, a marker indicative of proinflammatory microglia, was examined. Co-localization of CD68 with IBA1 staining (Fig. 2C, D, and F) revealed a significantly higher CD68/IBA1 ratio in TG mice compared to WT mice. Treatment with the DRD2 receptor agonist quinpirole reduced this ratio (n = 4 per group; two-way ANOVA: WT *vs* TG, P = 0.0013; WT *vs* WT + Quin, P = 0.9562; TG *vs* TG + Quin, P = 0.0293; TG + Quin *vs* WT, P = 0.4633). Collectively, these findings indicate that the hR1441G mutation polarizes microglia, altering both their distribution and functional state in the dorsal striatum. DRD2 receptor activation mitigates LRRK2 hR1441G-induced microglial proinflammatory responses.

**Figure 2.**
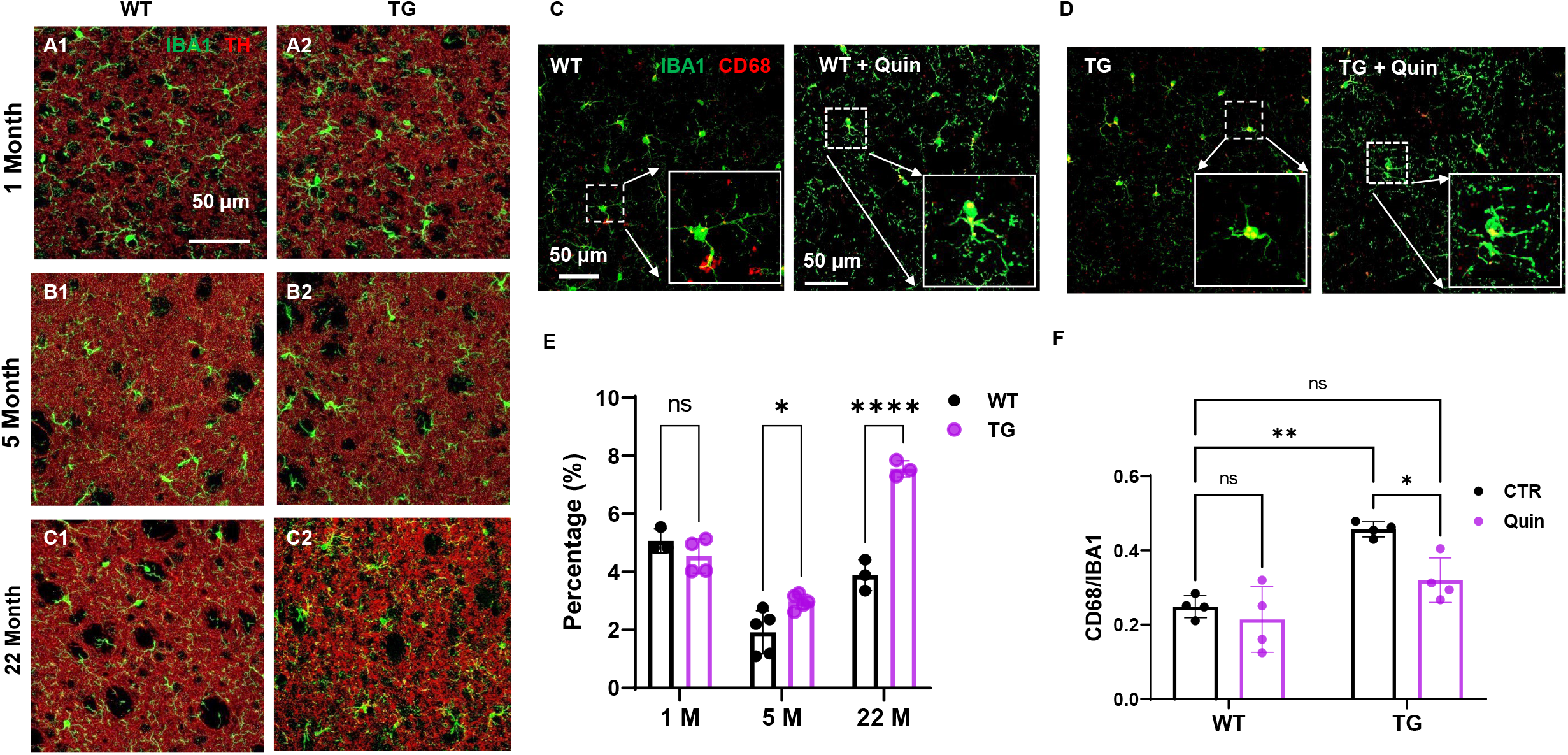
Microglia distribution and polarization in the LRRK2 hR1441G animal model normalized by DRD2 receptor agonist quinpirole. **(A1-A2, B1-B2 and C1-C2)** Visualization of the distribution of microglia within the dorsal striatum at 1, 5 and 22 months of age between hR1441G TG and WT mice. **(C and D)** Co-localization of IBA1 and CD68 markers in hR1441G TG and WT mice, respectively. Inset, magnification of the region of interest (ROI1 and ROI2). **(E)** Quantification of the percentage of microglia area per unit of the dorsal striatum (mm^2^). No significant difference of microglia coverage area in the 1-month-old mice between WT and TG mice was observed; at 5 months, WT mice displayed significantly higher microglial area compared to TG; This difference remained significant in 22-month-old mice (WT, N = 3, TG, N = 4, Two-way ANOVA, WT *vs* TG, 1-month-old, P value 0.4893; 5-month-old, P value 0.0149; 22-month-old, P value < 0.0001). **(F)** Quantification of CD68-positive area within microglia. Co-localization of IBA1 and CD68 revealed that CD68 area in TG microglia was significantly greater than in WT microglia and quinpirole treatment decreases the ratio of CD68/IBA1 (4 animals each group, two-way ANOVA, WT *vs* TG, P value 0.0013, WT *vs* WT + Quin, P value 0.9562, TG *vs* TG + Quin, P value 0.0293, TG + Quin. *vs* WT, P value 0.4633).

### LRRK2 hR1441G mutation decreased the response of microglia to focal injury in the dorsal striatum of acute brain slices

We sought to determine whether microglial polarization impacts microglial function, specifically response capability and dynamics, within the dorsal striatum. Acute coronal brain slices were prepared from CX3CR1-GFP mice. The dorsal striatum was imaged using 2PLSM. To directly assess the influence of the hR1441G mutation on microglial responses to external stimuli, a focal laser ablation (15-20 µm diameter) was performed 30 µm below the surface of the brain slice in the dSTR using an infrared laser beam at 890 nm, which employs a non-linear process to confine the injury to the focal center.

As shown in Fig. 3, microglia in proximity to the injury site exhibited migratory behavior, extending the processes toward the damaged area. Microglial located further away also responded but did not reach the injury site. Notably, the tips of the processes of these cells, located near the injury, appeared bulbous and slightly enlarged, extending toward the damage (Fig. 3A1-A2, B1-B2, C1-C2, and Supplementary Video 1). Approximately 10 minutes post-laser injury, the processes of adjacent microglia reached the damage site, forming a spherical containment around the lesion (Supplementary Fig. 4), while their cell bodies remained stationary. Compared to WT mice, the response of microglia in TG mice were both significantly delayed and reduced in magnitude. Our previous studies have shown decreased dopamine release in the dorsal striatum of hR1441G TG mice.^19,20^ To examine whether D2R was involved in microglial dysfunction within the dSTR, brain slices were treated with the D2R agonist quinpirole. This treatment significantly enhanced the response of microglia to injury (Fig. 3G).

**Figure 3.**
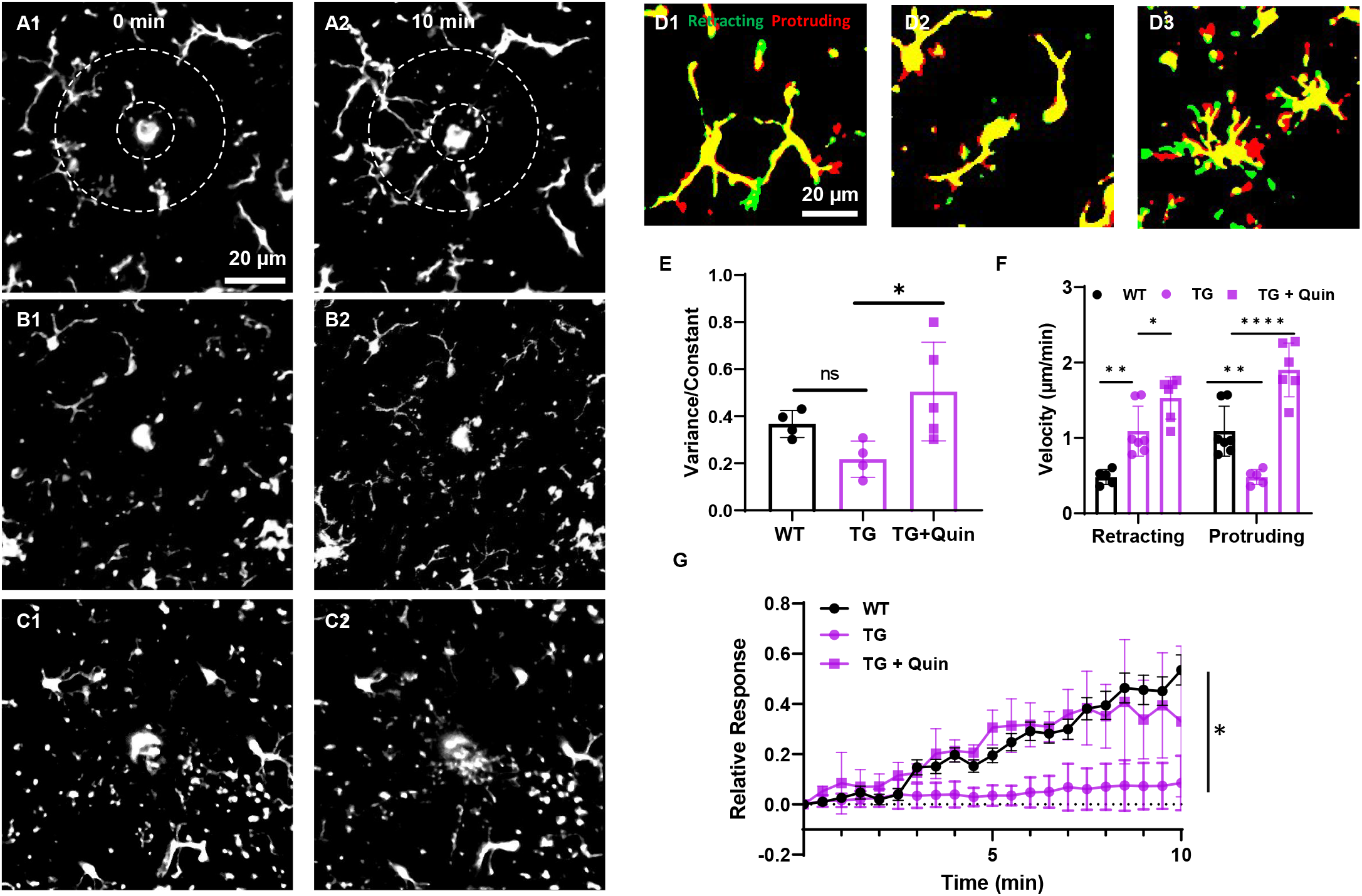
The D2 receptor agonist, quinpirole, ameliorates microglial dysfunction in the dorsal striatum of hR1441G Tg mice. **(A1-A2)** representative images depicting microglial response to laser ablation in WT at time 0- and 10-minutes post-ablation. **(B1-B2)** representative images of microglial response to laser ablation in TG mice at 0- and 10-minutes post-ablation. **(C1-C2)** representative images of microglial response to laser ablation in TG mice at 0 and 10 minutes following quinpirole treatment. **(D1-D3)** representative image of microglial dynamics between 0 and 5 minutes in WT, TG and TG + Quin treatment. **(E)** Quantification of variance region-to-constant region ratio of microglial between 0 and 5 minutes at the different condition, A significant difference was observed in this ratio between WT and TG and quinpirole treatment increased the variance-to-constant region ratio in TG mice (3 pairs, Mann-Whitney test, P value WT *vs* TG 0.0571, P value TG *vs* TG + Quin 0.0317). **(F)** Significant differences in both protrusion and retraction velocities were found between WT and TG, with quinpirole treatment significantly enhancing both in TG mice (4 pairs, two-way ANOVA, retracting: WT *vs* TG, P value 0.0019; TG *vs* TG + Quin, P value 0.0173; Protruding: WT *vs* TG, P value 0.0018; TG *vs* TG + Quin, P value < 0.0001). **(G)** Quinpirole treatment improves the response to microglia to laser ablation in TG mice.

We then determined whether this diminished response was related to the dynamics of microglia. We examined microglial protrusion and retraction under various conditions: WT, TG, and TG with quinpirole treatment (Fig. 3D1-D3 and Supplementary Video 2). The dynamics of microglia were monitored for 10 minutes. Between 0 and 5 minutes, the regions of protrusion and retraction in TG mice were less than those in WT mice (WT *vs* TG, Mann-Whitney test, P = 0.0571), but quinpirole treatment significantly enhanced microglial dynamics (Fig. 3E, Mann-Whitney test, TG + Quin *vs* TG, P = 0.0317). We further quantified the velocities of microglial protrusion and retraction in WT, TG, and TG with quinpirole treatment. Microglia in TG mice exhibited slower protrusion, but faster retraction compared to WT littermates (Fig. 3F, WT *vs* TG, 4 pairs, protrusion P = 0.0018; retraction P = 0.0019). Quinpirole treatment significantly increased the velocities of both protrusion and retraction (Fig. 3F, TG *vs* TG + Quin, retraction P = 0.0173; protrusion P < 0.0001). Thus, these findings support that quinpirole improved microglial responses to laser ablation.

### LRRK2 hR1441G mutation decreased the response of microglia to focal brain injury *in vivo*

To mimic physiological conditions and minimize the impact of mechanical slicing injury on microglial function in brain slices, we employed an intravital two-photon microscopy system to study microglial dynamics and function *in vivo*. Imaging was conducted through PoRTS procedure over the somatosensory cortex.

As illustrated in Fig. 4, following laser ablation at a depth of 100 μm below the surface, the enlarged tips of microglial processes extended toward the site of injury (Fig. 4A1-A2, B1-B2, C1-C2, and Supplementary Video 3). Approximately 20 minutes post-injury, the processes of nearby microglia reached the damage site, forming a spherical containment, while the cell bodies remained stationary (Supplementary Fig. 5). These findings closely resemble the phenomena in the dorsal striatum of acute brain slices. Furthermore, compared to WT littermates, the response of microglia in TG mice was both slower and less pronounced; similarly, quinpirole treatment enhanced the microglial response to injury (Fig. 4G).

**Figure 4.**
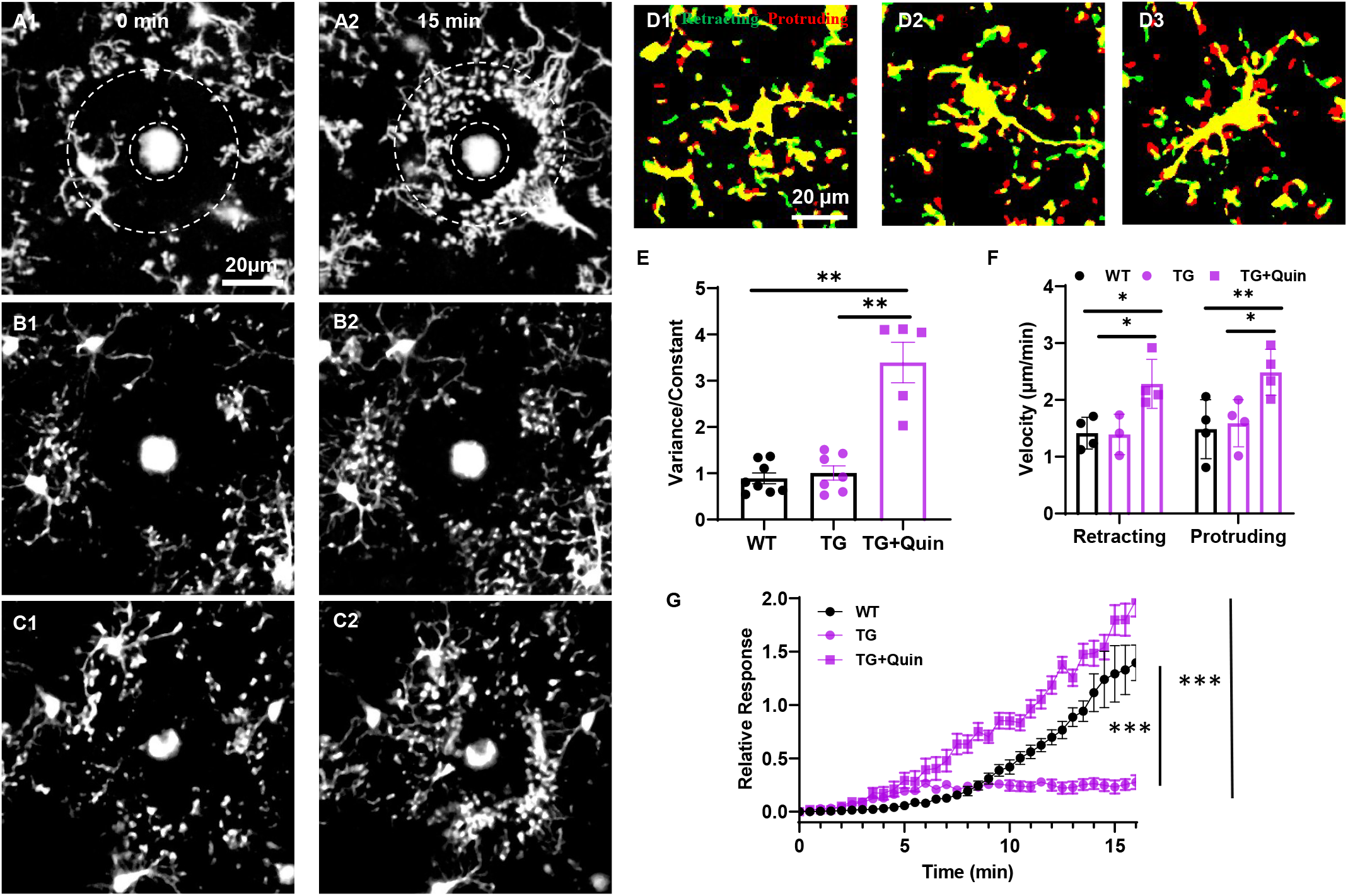
The D2 receptor agonist, quinpirole, ameliorates the deficits in cortical microglial response to external stimuli in the hR144G Tg mice *in vivo*. **(A1-A2)** representative images of microglial response to laser ablation in WT at 0 minute and 15 minutes. **(B1-B2)** representative images of microglial response to laser ablation in TG at 0 and 15 minutes. **(C1-C2)** representative images of microglial response to laser ablation in TG mice at 0 and 15 minutes following quinpirole treatment. **(D1-D3)** representative image illustrating microglial dynamics in WT, TG and TG + Quinpirole treatment between 0 and 5 minutes. **(E)** Quantification of the variance-to-constant region ratio of microglia between 0 and 5 minutes at the different condition. No significant differences between WT and TG, but quinpirole significantly increased this ratio in TG mice (3 pairs, Mann-Whitney test, WT *vs* TG, P value 0.6126; TG + Quin *vs* TG, P value 0.0025; TG + Quin, P value 0.0016). **(F)** Quantification of the velocities of microglial protrusion and retraction under different conditions. No significant differences in velocities were found between WT and TG. However, quinpirole increased both protrusion and retraction velocities in TG mice (3 pairs, two-way ANOVA, retracting: WT *vs* TG, P value 0.9998; WT *vs* TG + Quin, P value 0.0246; TG *vs* TG + Quin, P value 0.0338; protruding: WT *vs* TG, P value 0.9786; WT *vs* TG+ Quin, P value 0.0093; TG *vs* TG + Quin, P value 0.0202). **(G)** Quinpirole treatment enhances the microglial response to laser ablation in LRRK2 R1441G TG mice.

To explore whether this effect was related to microglial dynamics, we examined the retraction and protrusion of microglia under various conditions: WT, TG, and TG + Quin (Fig. 4D1-D3 and Supplementary Video 4). Microglial dynamics were monitored for at least 15 minutes. The areas of protrusion and retraction relative to a constant area remained consistent between 0 and 5 minutes in both WT and TG mice. Quinpirole treatment significantly increased the area of variance in microglial dynamics compared to TG (Fig. 4E, Mann-Whitney test, WT *vs* TG, P = 0.6126; TG + Quin *vs* TG, P = 0.0025). The velocities of retraction and protrusion were similar between WT and TG. Notably, quinpirole treatment significantly increased the velocities of both protrusion and retraction (Fig. 4F, three pairs, two-way ANOVA, WT *vs* TG, protrusion P = 0.9786; retraction P = 0.9998; TG *vs* TG + Quin, retraction P = 0.0338; protrusion P = 0.0202).

## Discussion

Emerging evidence highlights the immune system’s critical role in PD pathogenesis, particularly through microglial dysfunction. LRRK2, the leading genetic contributor to both familial and sporadic PD, has been implicated in driving this connection. Here, we provide the first evidence of how the pathogenic LRRK2-R1441G mutation disrupts microglial function *in vivo*, linking it directly to PD progression. Specifically, the R1441G mutation was found to impair microglial dynamic capacity and attenuate their responses to external stimuli. This dysfunction likely results in diminished microglial phagocytic activity toward aggregated proteins, promoting neuroinflammation and exacerbating disease pathology. These findings establish a critical connection between LRRK2 mutations, impaired microglial activity, and neurodegenerative processes in PD. Notably, treatment with the D2 receptor agonist quinpirole not only alleviates these deficits but also enhanced microglial dynamics and improved their response to external stimuli. These findings suggest that quinpirole may mitigate neuroinflammation associated with the hR1441G induced microglial polarization in the context of PD.

Microglia, long-lived cells tightly regulated throughout postnatal CNS development, increase in number during the first fourteen postnatal days, followed by a decline in the third postnatal week, resulting in a 50% reduction by six weeks of age.^17,18^ Single-cell imaging studies have indicated that the median lifespan of neocortical microglia exceeds 15 months, with approximately half surviving throughout the lifespan of the mouse. Additionally, a threefold increase in microglia numbers has been observed in the neocortex of aged APPPS1 mice.^21,22^ Our findings align with these studies, revealing a decrease in microglial numbers from 1 month old to young adulthood, which was less pronounced in TG mice compared to WT controls. This suggests that hR1441G mutation may drive the proliferation of microglia and the differentiation of monocytes in the brain. Furthermore, we found that microglial numbers in aged TG mice exceeded those in WT mice and even surpassed counts observed in 1-month-old mice.

Previous reports have indicated that quinpirole administration reduces neuroinflammation and mitigates brain injury in models of intracerebral hemorrhage (ICH) and traumatic brain injury (TBI) ^11,23^. Similarly, in the LRRK2 hR1441G TG mice, we observed D2 receptor modulation meliorate microglia dysfunction by restoring mobility and responsiveness. Long-term quinpirole treatment intriguingly attenuated microglial responsiveness to external stimuli in the dorsal striatum.

At current, two-photon laser scanning microscopy (2PLSM) faces challenges in imaging regions deeper than 1 mm beneath the cortical surface, such as the hippocampus and striatum.^24^ Nonetheless, *in vivo* 2PLSM has been employed to functionally image hippocampal place cells^25^ and investigate striatal circuit functions at cellular resolution.^26^ However, these studies often require surgical removal of cortical tissue, which can trigger neuroinflammation, including microglial activation. Patients with PD exhibit sensory and motor dysfunction, including impaired sensory processing, which can recover with dopaminergic therapy and deep brain stimulation.^27^ Therefore, in this study, we selected the somatosensory cortex, rather than the striatum, for investigating microglial dynamics and functions *in vivo*. To minimize such inflammation and maintain brain integrity, we implemented a procedure known as PoRTS, a skull-thinning technique,^14^ to minimize tissue damage. Future advancements in depth penetration for two-photon imaging will be crucial for monitoring microglial dynamics and function within the striatum.

## Abbreviations

D2R=: dopamine D2 receptor
dSTR=: the dorsal striatum
LRRK2=: Leucine-rich repeat kinase 2
PD=: Parkinson’s disease
SNc=: substantia nigra pars compacta

## Data availability

Data is available from the corresponding authors on request.

## Supplementary material

Supplementary material is available online.

