## Supplementary materials for "Quinpirole ameliorates the dysfunction of microglia in human LRRK2-R1441G transgenic mice"

### Supplementary Data

#### Supplementary methods

##### Animals and brain slice preparation

BAC LRRK2 (hR1441G) transgenic (Tg) mice (stock #009604, The Jackson Laboratory) were obtained from Chenjian Li's laboratory at Weill Medical College of Cornell University and maintained on Taconic FVB/N background. CX3CR1-GFP transgenic mice (Jackson Laboratory, strain #005582) were backcrossed onto an FVB/N background for eight generations, and then crossed with hR1441G Tg mice to create hR1441G-Tg/CX3CR1-GFP$^+$ and hR1441G-WT (nTg)/ CX3CR1-GFP$^+$ mice. All mice were on FVB/N mouse background (Taconic). The mice were housed in a temperature and humidity-controlled condition with 12/12 hours light/dark cycle and allowed free access to food and water in a specific pathogen-free facility. All animal procedures were conducted in accordance with the National Institutes of Health guidelines and were approved by the Institutional Animal Care and Use Committee (IACUC) of Thomas Jefferson University and The University of Georgia.

For the preparation of acute brain slices, mice were anesthetized with a ketamine/xylazine mixture (100 mg/10 mg/kg), followed by transcardial perfusion with ice-cold, oxygenated cutting solution (in mM: 125 NaCl, 2.5 KCl, 26 NaHCO3, 3.7 MgSO4, 0.3 KH2PO4, 10 glucose, pH 7.4). After perfusion, the mice were decapitated, and their brains were immediately extracted. Coronal brain sections were then obtained at a thickness of 300 μm using a vibratome (VT1000S, Leica Microsystems, Germany) and were subsequently incubated at 34°C in oxygenated artificial cerebrospinal fluid (aCSF) (in mM: 125 NaCl, 2.5 KCl, 26 NaHCO3, 2.4 CaCl2, 1.3 MgSO4, 0.3 KH2PO4, 10 glucose, 2 HEPES, pH 7.4) for 30 minutes before being transferred to room temperature.

#### Tissue preparation and Immunofluorescence staining

Animals were anesthetized via intraperitoneal injection of a Ketamine/Xylazine mixture (100 mg/10 mg/kg) and subsequently underwent transcardial perfusion with 30 mL of phosphate-buffered saline (PBS), followed by 30 mL of ice-cold 4% paraformaldehyde (PFA) fixative, prepared in PBS (pH 7.4), utilizing a perfusion pump. The brain was then extracted and post-fixed in PFA for 24 hours, after which it was submerged in 30% sucrose in PBS for 3 days. The brain was embedded in optimal cutting temperature (OCT) compound and sectioned coronally at 30μm using a Leica cryostat (Leica Microsystems, Germany). Serial sections of the striatum (STR) and substantia nigra (SN) were collected into twelve wells, with each well containing ten sections. The sections were cryopreserved in an antifreeze buffer composed of 30% glycerol and 30% ethylene glycol in PBS at −20 °C.

For immunohistochemical analysis, sections were blocked in PBS containing 0.2% Triton X-100 (PBST) and 5% donkey serum for 1 hour at room temperature. They were then incubated with primary antibodies, diluted in the blocking solution, at 4 °C for 36 hours. Following three rinses with PBST, the sections were incubated with appropriate fluorescent probe-conjugated secondary antibodies (DyLight 594-conjugated and DyLight 488-conjugated, 1:500; Life Technologies) for 45 minutes at room temperature with gentle agitation. The primary antibodies included anti-tyrosine hydroxylase (TH) (Millipore, MAB5280, 1:500), anti-IBA1 (Wako, AB_839504, 1:500), and anti-CD68 (Bio-Rad, AB_322219, 1:500). After three additional washes in PBST and one wash in PBS, sections were mounted using Dako Fluorescence Mounting Medium (Dako North America, Inc., S3023). Images were captured using a Zeiss LSM 900 confocal microscope.

#### Cranial window Surgery: polished and reinforced thinned skull (PoRTS) procedure

Three to four male pairs of hR1441G-Tg/CX3CR1-GFP$^+$ and hR1441G-WT (nTg)/ CX3CR1-GFP$^+$ mice (6–7 months old) were anesthetized via inhalation of isoflurane (4% for induction; 2% for surgical procedures) and secured in a custom-designed stereotactic apparatus. The body temperature was monitored using a rectal probe and maintained at 37.0°C with a heating blanket (Homeothermic Blanket System, Harvard Apparatus, Holliston, Massachusetts). Following the removal of fur from the skull, the parietal bone was exposed, and a region of 1–1.5 mm in diameter over the somatosensory cortex was carefully thinned to a depth of 20–40 μm using a high-speed surgical drill. Cold saline was applied to dissipate frictional heat and prevent damage to the underlying cortex.[14] Experiments were conducted only if physiological parameters remained within normal ranges.

#### Two-photon laser ablation

Laser-induced injury was generated by directing a two-photon laser beam to a confined region at a depth of 70–150 μm through a thinned, intact skull for *in vivo* imaging, or at a depth of 30–50 μm below the surface of brain slices in the dorsal striatum for *ex vivo* imaging. The two-photon laser was tuned to a wavelength of 890 nm, with laser power maintained between 100-150 mW. The beam was focused on the targeted area for approximately 3–5 seconds to induce a localized injury, which was confined to a region with a diameter of approximately 20–30 μm.[13]

#### 2PLSM: Imaging of microglia in acute brain slices and *in vivo*

The upright laser scanning microscope (BX61WI, Olympus), coupled with a Ti: sapphire pulsed laser system (80 MHz repetition rate, <100 fs pulse width, Coherent Inc.), and operated using Prairie View 5.3 software (Bruker), was utilized for two-photon imaging (Fig. S1). Objective lenses with 20x magnification (NA 1.0; WD 2.0 mm, Olympus) and 40x water immersion (NA 0.8; WD 3.3 mm, Olympus) were selectively employed for both *ex vivo* and *in vivo* imaging.

**Acute Brain Slices:** Acute coronal brain slices from mice expressing CX3CR1-GFP were positioned in a recording chamber with continuous perfusion of oxygenated artificial cerebrospinal fluid (aCSF) at physiological temperatures (34–35°C). The dorsal striatum was designated as the region of interest. GFP fluorescence (500–550 nm) was isolated using a 525/50 nm filter and detected via a non-descanned photomultiplier tube (PMT). Laser intensity and PMT gain were held consistent throughout the time-lapse imaging. Imaging depth was restricted to 30 μm below the surface of the section, with time-series scanning (20–60 frames) capturing stacks of 20–30 image planes at 1 μm axial intervals, using a 4-μs dwell time and a rate of 30 s per stack frame, excluding any drifted frames. The average laser power for imaging was kept below 50 mW.

***In Vivo* Imaging:** Animals that underwent PoRTS surgery were positioned under the two-photon laser scanning microscope (2PLSM) for microglial imaging. Time-lapse imaging of small cortical sub-volumes (20–30 image planes with 1 μm axial spacing) was performed for a minimum of 15 minutes to monitor the dynamics of microglia expressing EGFP in the cerebral cortex. The interval between stack sequences was maintained at 25–30 seconds, with PMT settings (including gain and offset) and laser excitation power kept constant throughout the time-lapse imaging. Imaging depth was maintained at 70 μm below the surface. Experiments exhibiting unintentional photoactivation were excluded from analysis.

To investigate the effects of the D2 receptor agonist quinpirole on microglial dynamics and function, brain slices were incubated in aCSF solution containing quinpirole (3 μM, Cat. #1061, Tocris) for 30 minutes, followed by perfusion with aCSF containing 1 μM quinpirole during imaging. *In vivo* imaging of microglia was conducted following a single intraperitoneal injection of quinpirole (0.5 mg/kg).[15]

#### Data analysis

Images were processed using the open-source software Fiji (NIH) and commercial software Matlab (Version 8.5.0 R2015a, MathWorks). Maximum intensity projection (MIP) of *z*-stacks was employed to generate two-dimensional (2D) representations of three-dimensional (3D) structures. All analyses of time-lapse movies were conducted using MIP of sequentially acquired image stacks. To correct for lateral drift along the x and y axes, image registration was performed to achieve intensity-based alignment across different time points. To investigate the dynamics of microglial protrusions and retractions, 3D image stacks were captured at all time points above and below the microglia of interest. The length of microglial processes and the velocity of length changes were assessed from MIP images using Fiji software.[16]

As described by Davalos[13], to quantify the microglial response to laser ablation, two regions surrounding the ablation site were defined: the outer area (Y) with a radius of 70 μm and the inner area (X) with a radius of 35 μm. The number of microglial processes migrating from the outer area into the inner area was measured over time. Threshold was set for each image to produce a binary representation, and the number of white pixels in the inner area over time (Rx(t)) was quantified as an indicator of microglial response to laser ablation. To account for variations in the distribution and number of microglia within the outer area, the microglial response was calculated relative to the number of processes in the outer area immediately after ablation (Ry(0)). The microglial response at time point t (R(t)) was computed using the following equation: $R(t) = [Rx(t) – Rx(0)] / Ry(0)$, Rx(t), where Rx(t) represents the number of white pixels at time point t, while Rx(0) and Ry(0) represent the number of white pixels at the initial time point within the inner and outer areas, respectively.

To evaluate the level of microglial polarization, double immunostaining for IBA1 and CD68 was performed on fixed brain slices to assess co-localization of these markers. Maximum intensity projections (MIPs) of 15 image stacks, including both IBA1/CD68 and TH/IBA1, were generated. Three fields from the dorsal striatum of each animal were randomly selected for quantitative analysis of microglia. Co-localization of IBA1 and CD68 was analyzed to quantify CD68 fluorescence within microglia, with the total area of internalized CD68 calculated. Microglial polarization was determined using the following formula: polarization ratio = area of internalized CD68 / area of the microglial cell.

#### Digital spatial profiling

**FFPE Tissue Preparation:** Formalin-fixed paraffin-embedded (FFPE) tissues were prepared as follows: mice were anesthetized, and brains were fixed via transcardial perfusion with 10% neutral buffered formalin (Sigma-Aldrich). Brains were subsequently removed, post-fixed in the same formalin solution for 24 hours at 4°C and immersed in 70% ethanol for at least 1 hour. Tissues were processed and embedded in paraffin using an automated tissue processor (Tissue-Tek VIP, Sakura Finetek USA Inc.) following the programmed protocol: 95% ethanol for 75minutes, 100% ethanol for 75minutes, 100% ethanol for 90 minutes, 100% xylene for 1h, 100% xylene for 90 minutes, 100% xylene for 90 minutes, paraffin for 90 minutes at 58 °C, and paraffin for 2 hours at 58°C. FFPE tissues were then immersed in molten paraffin at 60°C in a metal mold to form FFPE blocks. These blocks were cooled at 4°C and stored at the same temperature until further use.

**Tissue Sectioning and Slide Preparation:** The midbrains of FFPE mouse brains were coronally sectioned at 5 µm using a rotary microtome. Sections containing the substantia nigra (SN) and ventral tegmental area (VTA) were mounted on Leica BOND Plus microscope slides, with each slide holding 6 sections—3 from WT mice and 3 from Tg mice. The slides were submitted to the Flow Cytometry and Human Immune Monitoring Core at Thomas Jefferson University for RNA profiling via the GeoMx RNA assay (Nanostring Technologies, Seattle, WA, USA).

**ROI and AOI Selection:** Tissue sections were stained with fluorescent morphology markers and nuclear counterstain for the identification of regions of interest (ROIs) and areas of interest (AOIs). The morphology markers included an anti-tyrosine hydroxylase (TH) antibody conjugated with Alexa 488 (dopaminergic neuron marker, Sigma #MAB5280X), and an anti-Iba1 antibody conjugated with Alexa 647 (microglia marker, Millipore #MABN92-AF647). Syto-13 was used as a nuclear marker. For each tissue section, 2-3 ROIs were selected within the SN or VTA, and each ROI was subdivided into 1-3 AOIs representing TH+ or Iba1+ cell populations. Each AOI contained a minimum of 20 cells.

**Quality Control:** Raw counts from the GeoMxTM Digital Spatial Profiling (DSP) were processed using GeoMxTM Software. Out of 95 segments collected from 6 mice (3 hR1441G Tg mice and 3 WT mice), 11 segments were excluded due to failure in quality control (QC) analysis. Following QC, 8021 out of 20175 gene targets in the GeoMx Mouse Whole Transcriptome Atlas passed QC and were retained for downstream analyses.

**Data Normalization and Analysis:** To address systematic variability between AOIs, GeoMxTM DSP raw counts were normalized to the 75th percentile of expression for each AOI, as described in the NanoString GeoMxTM Data Analysis User Manual. Differential expression analysis was performed using a negative binomial model, implemented in the DESeq2 package in R.

**Gene Set Enrichment Analysis (GSEA):** Gene Set Enrichment Analysis (GSEA) was carried out to identify dysregulated pathways in microglia between mice carrying hR1441G mutation and WT controls. GSEA was conducted using the ClusterProfiler package (version 4.12) in R, with gene sets ranging in size from 10 to 500 genes. Enrichment terms with p-values below 0.05 were considered significantly enriched. Additionally, GSEA was extended using gene set signatures from the Gene Ontology database, with normalized enrichment scores (NES) calculated to identify biological pathways linked to hR1441G mutation in microglia.

#### Supplementary Figure 1

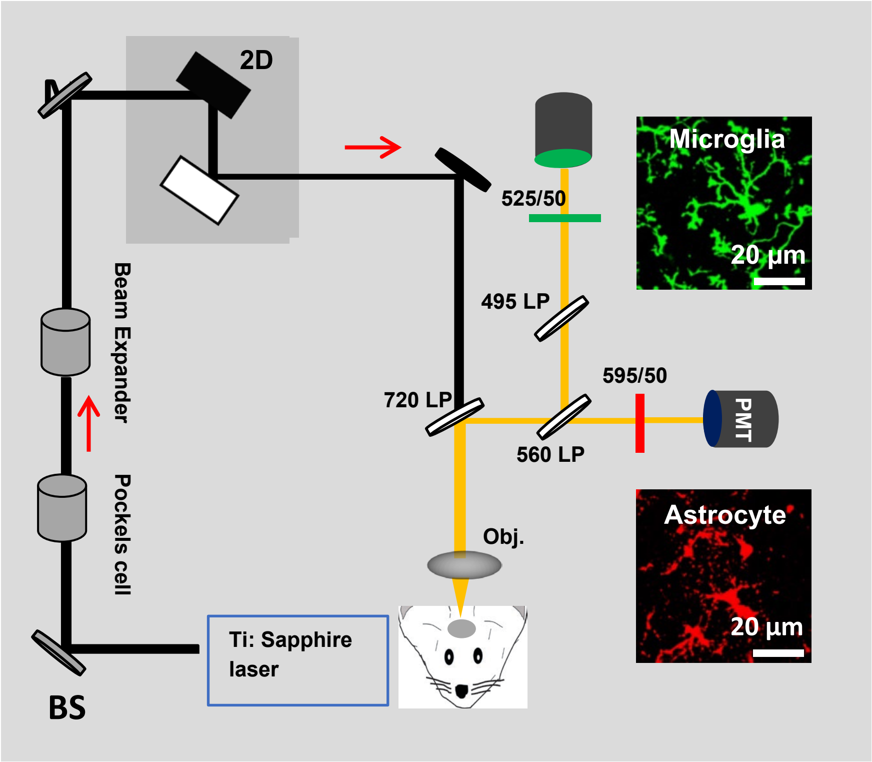


**Supplementary Figure 1. Schematic illustration of 2PLSM system**

#### Supplementary Figure 2

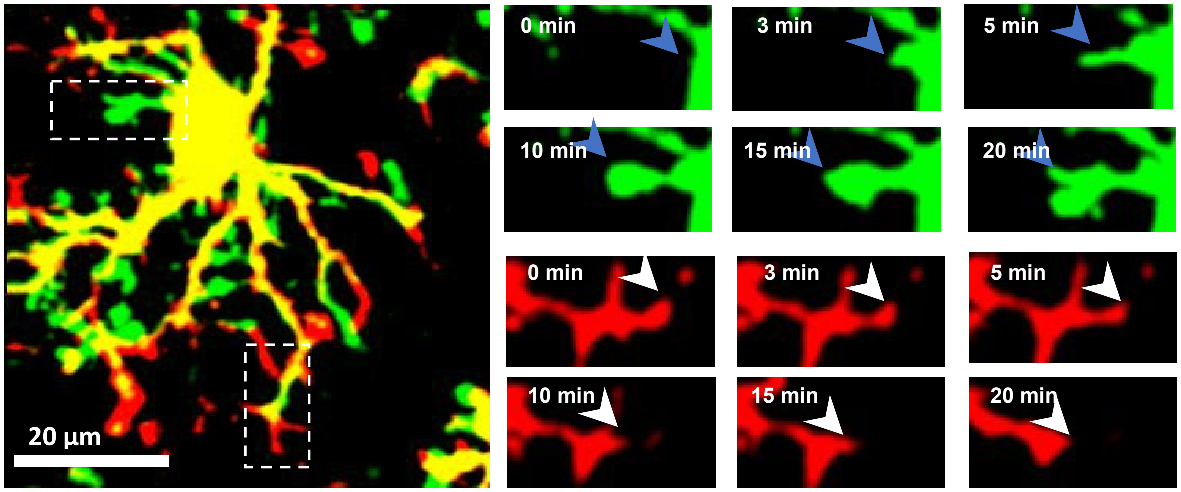


**Supplementary Figure 2. Velocity measurement of microglia dynamics, showing both protruding and retracting.**

**Supplementary Figure 3**

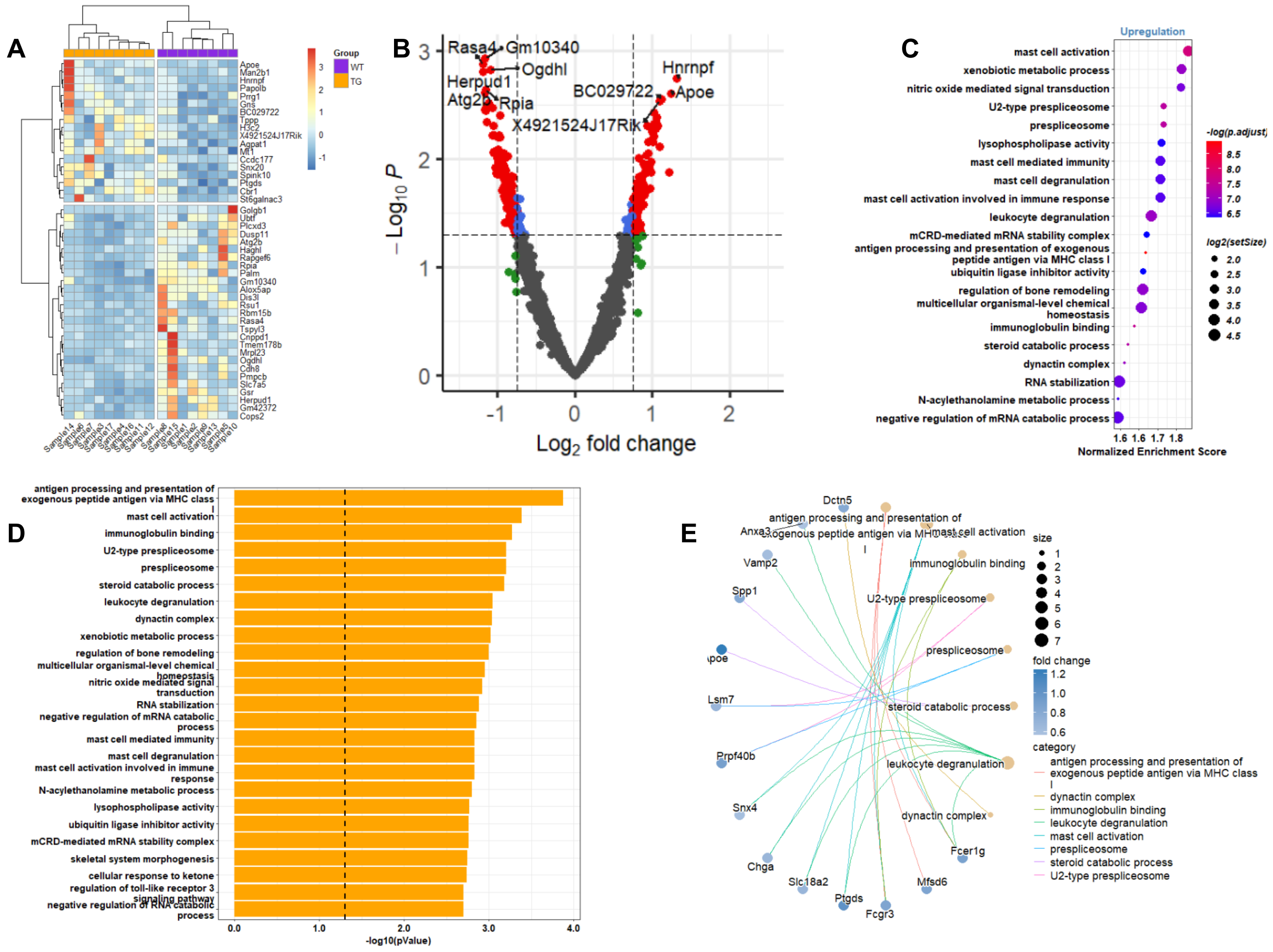

**Supplementary Figure 3. GeoMx digital spatial profiling (DSP) of microglia in the substantia nigra pars compacta (SNc).**

**(A)** Heatmap displaying differentially expressed genes. **(B)** Volcano plot illustrates the distribution of differentially expressed genes. **(C)** Gene ontology enrichment analysis of upregulated genes, with terms colored by p-values and sized according to gene set size. **(D)** Bar plot showing the top 20 enriched terms, ordered by P-value. **(E)** Gene network visualization of enriched terms.

**Supplementary Figure 4**

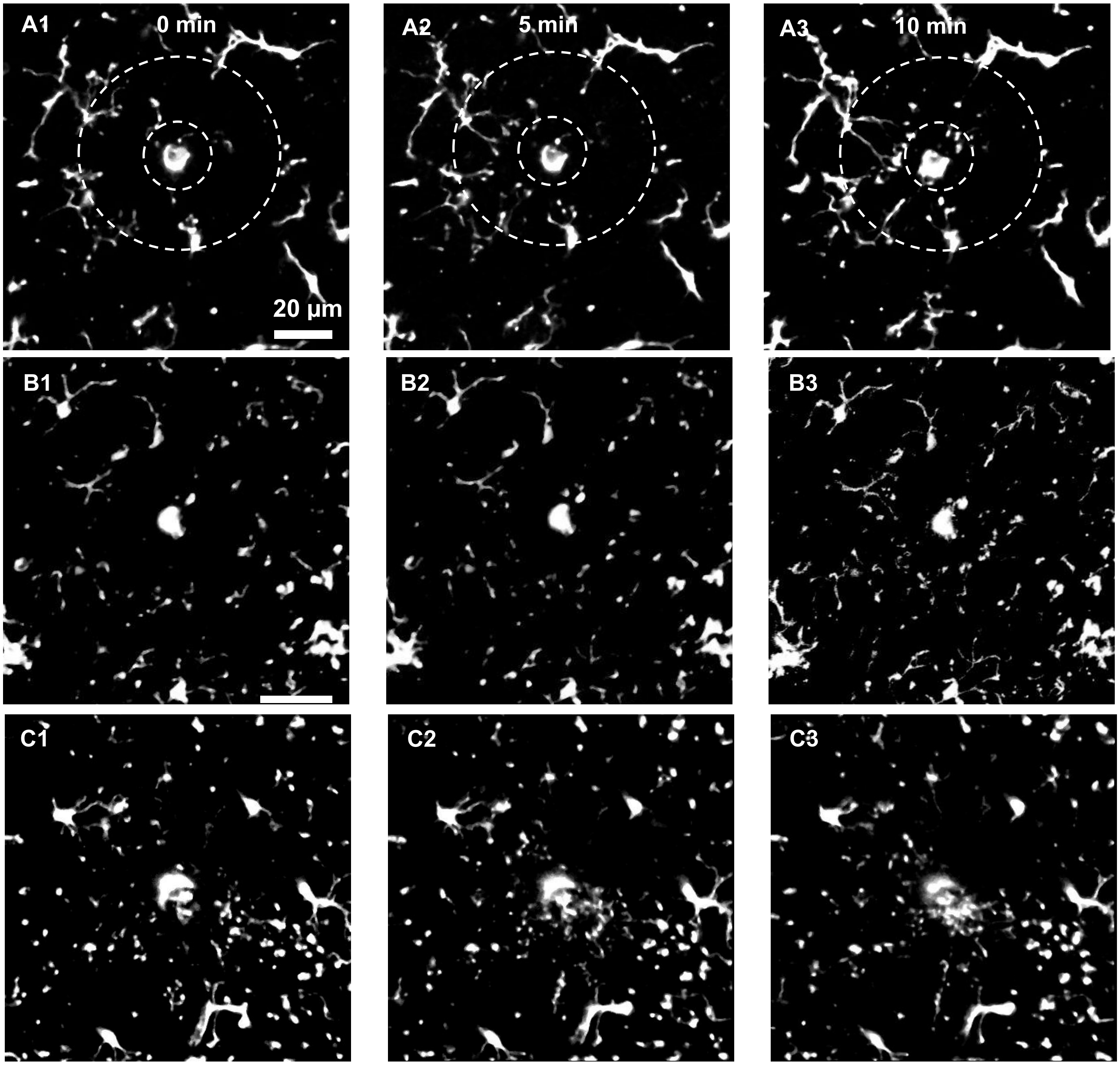


**Supplementary Figure 4. Visualization of the response of the microglia in the dorsal striatum of acute brain slice from hR1441G Tg mice and WT controls to the external stimulus: WT, LRRK2 and LRRK2 + quinpirole.**

**(A1-A3)** representative images of microglial response to laser ablation in WT at time 0, 5 and 10 minutes. **(B1-B3)** representative images of microglial response to laser ablation hR1441G at time 0, 5 and 10 minutes. **(C1-C3)** representative images of microglial response to laser ablation in hR1441G at time 0, 5 and 10 minutes after quinpirole treatment.

**Supplementary Figure 5**

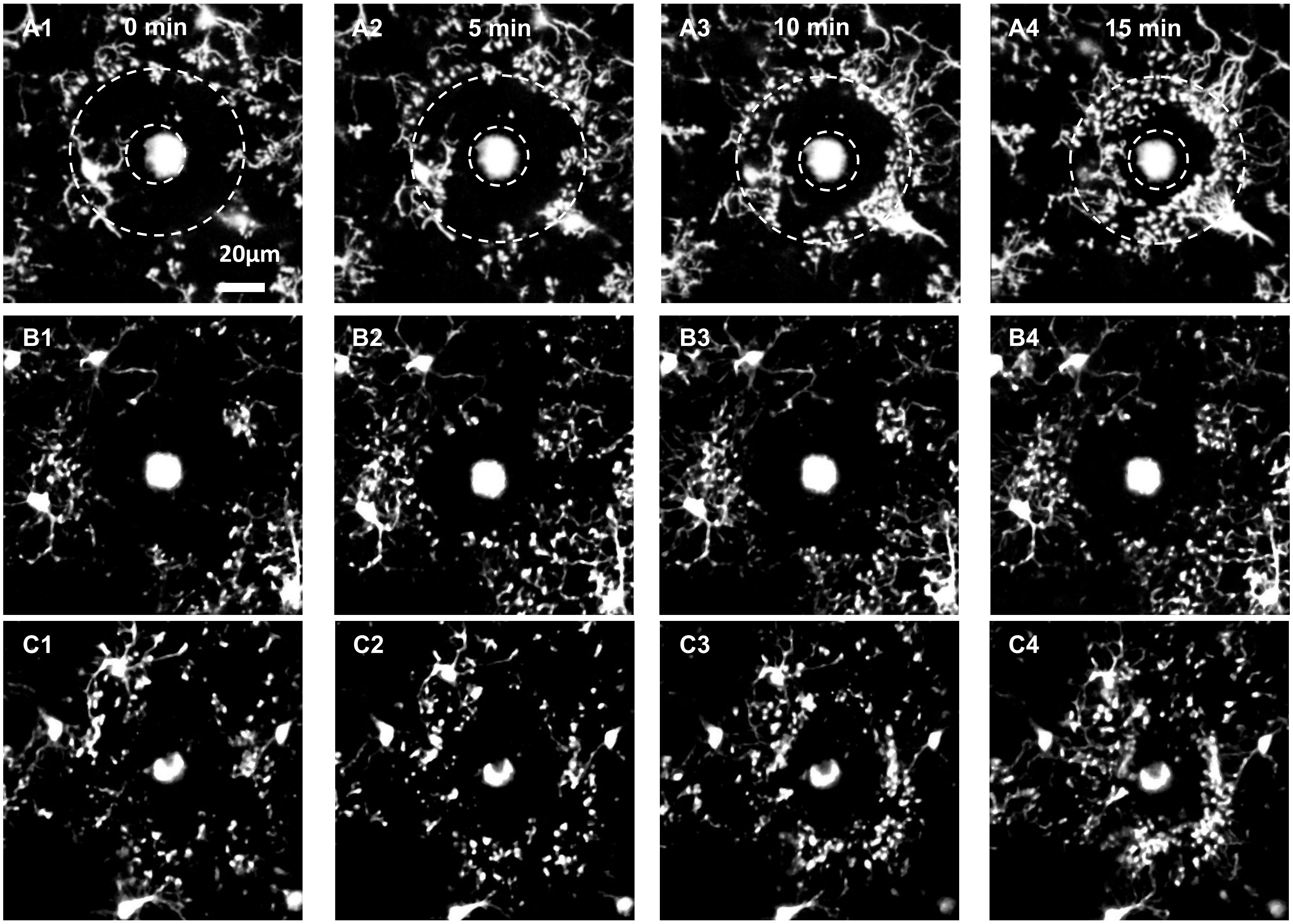


**Supplementary Figure 5**. ***In vivo* visualization of microglial responses in the somatosensory cortex from hR1441G Tg mice and WT controls to external stimuli: WT, hR1441G, hR1441G treated with quinpirole.**

**(A1–A4)** Representative images of microglial responses to laser ablation in WT at 0, 5, 10, and 15 minutes. **(B1–B4)** Representative images of microglial responses to laser ablation in hR1441G at 0, 5, 10, and 15 minutes. **(C1–C4)** Representative images of microglial responses to laser ablation in hR1441G at 0, 5, 10, and 15 minutes following quinpirole treatment.

#### Supplementary Video Legends

**Supplementary Video 1**: Microglial responses to external stimuli in the dorsal striatum of acute brain slices from hR1441G Tg mice and WT controls: WT, hR1441G, and hLR1441G with quinpirole treatment.

**Supplementary Video 2**: Microglial dynamics in the dorsal striatum of acute brain slices from hR1441G Tg mice and WT controls: WT, LRRK2 hR1441G, and LRRK2 hLR1441G with quinpirole treatment.

**Supplementary Video 3**: *In vivo* visualization of microglial responses to external stimuli in motor cortex from hR1441G Tg mice and WT controls: WT, hR1441G, and hLR1441G with quinpirole treatment.

**Supplementary Video 4**: *In vivo* visualization of microglial dynamics in the **somatosensory** cortex from hR1441G Tg mice and WT controls: WT, hR1441G, and hLR1441G with quinpirole treatment.
